# Acute hypoxia induces transient olfactory dysfunction through olfactory epithelial degeneration and bulbar mitochondrial stress in zebrafish

**DOI:** 10.64898/2026.03.23.713737

**Authors:** Skylar L. DeWitt-Batt, Kate E. DeMann, Cameron J. Houck, Cassidy L. Larson, Luke A. Horsburgh, Evan A. Thomas, Liz Sanchez, Erika Calvo-Ochoa

**Affiliations:** Biology Department and Neuroscience Program, Hope College, Holland, MI, USA; Department of Neuroscience, University of Rochester Medical Center, Rochester, New York

**Keywords:** Olfactory system, zebrafish, neurogenesis, olfactory dysfunction, neuroinflammation, hypoxia, ischemia, neurodegeneration

## Abstract

Hypoxic-ischemic injury is a major cause of olfactory dysfunction, yet the cellular and morphological mechanisms underlying this sensory loss remain poorly understood. Here, we investigated the structural, cellular, and functional effects of acute hypoxic exposure on the olfactory system of adult zebrafish (*Danio rerio*) of both sexes, a model organism with remarkable neuroregenerative capacity. Fish were subjected to 15 minutes of acute severe hypoxia (0.8 mg/L dissolved oxygen) and assessed at 1 and 5 days post-hypoxia (dph). We evaluated olfactory function by means of cadaverine-evoked aversive behavioral assays. Structural and morphological integrity and inflammation of the olfactory epithelium (OE) and olfactory bulb (OB) were characterized using immunohistochemistry, histological stainings, and a 2,3,5-triphenyltetrazolium chloride (TTC) colorimetric assay. Acute hypoxic exposure impaired olfactory-mediated behaviors without affecting locomotion or exploratory behavior. In the peripheral OE, hypoxia caused neurodegeneration, disruption of the nasal mucus layer, and robust leukocytic infiltration. We observed reduced mitochondrial dehydrogenase activity in the olfactory bulb (OB) along with reactive astrogliosis. Olfactory function recovered by 5 days, coinciding with full restoration of OE morphology, and supported by a strong proliferative response. These findings reveal a coordinated degenerative and regenerative response to hypoxia across the olfactory axis, with implications for understanding hypoxia-induced sensory loss and neural repair.

**SIGNIFICANCE:** This work addresses an important gap in knowledge regarding the mechanisms linking hypoxic insult and olfactory dysfunction. By using adult zebrafish, an extraordinarily regenerative vertebrate, it also provides insight into neuronal repair and regenerative processes supporting olfactory recovery. The novelty of our study resides in that, to our knowledge, there are no studies that provide a comprehensive characterization of the effects of hypoxia in the olfactory system across molecular, histological, and functional levels. These findings advance our understanding of hypoxia-induced sensory neurodegeneration and regeneration, and highlight the zebrafish olfactory system as a powerful model for investigating neural repair mechanisms relevant to hypoxic-ischemic brain injury.

## INTRODUCTION

Hypoxia, defined as insufficient oxygen in tissues to sustain physiological functions, poses a significant threat to the health of terrestrial and aquatic organisms. In clinical settings, hypoxia can arise from a wide range of conditions including ischemic stroke (Feng et al., 2020; Leu et al., 2019), traumatic brain injury (Scultetus et al., 2016; Yan et al., 2014), and sleep apnea (Binar & Gokgoz, 2021; Hernandez-Soto et al., 2021).

Neurons are highly metabolic cells that rely on oxidative phosphorylation to meet their extensive energetic demands, making them particularly vulnerable to oxygen deprivation (Busl & Greer, 2010). Even brief hypoxic exposure can compromise mitochondrial function, leading to neuronal damage and initiate cell death pathways (Douglas et al., 2010), leading to a broad spectrum of sensory and cognitive deficits (Hernandez-Soto et al., 2021; Y. Lee et al., 2018; Wang et al., 2022).

Although olfactory loss is a well-recognized consequence of hypoxic-ischemic brain injury (Bigdaj et al., 2018; Hernandez-Soto et al., 2021; Huppertz et al., 2018; Kim et al., 2023), the mechanisms underlying its onset and potential recovery are poorly understood. Addressing this gap necessitates an experimental model that provides the opportunity to investigate both degenerative and regenerative processes following hypoxic insult. The olfactory system of adult zebrafish offers such a uniquely tractable model, given its well-characterized regenerative and neurogenic capabilities (Calvo-Ochoa et al., 2019; Calvo-Ochoa et al., 2020) and the availability of olfactory-mediated behavioral assays that allow functional outcomes to be linked to morphological states (Calvo-Ochoa et al., 2024; Hentig & Byrd-Jacobs, 2016). These features enable the study of both degenerative and regenerative processes within the same system and across recovery (Vorhees et al., 2025; White et al., 2015)

The architecture of the zebrafish olfactory system is similar to that of mammals, and it comprises three important components across the peripheral and central nervous system (Byrd & Brunjes, 1995). The peripheral olfactory epithelium (OE), located in the olfactory cavity, contains a densely-packaged array of olfactory sensory neurons (OSNs) that detect odorants via specialized cilia that extend into the mucus layer lining the epithelial surface (Ahuja et al., 2013; Hansen & Zielinski, 2005; Yoshihara, 2009). These neurons transmit sensory signals through the olfactory nerve (ON), a bundle of axons traversing the ethmoid bone, to the centrally located olfactory bulb (OB) (Mori et al., 1999; Sato et al., 2005), where sensory signals are relayed to higher brain centers for further processing (Miyasaka et al., 2009).

The olfactory system is uniquely vulnerable to low-oxygen conditions for two reasons: first, the OE is directly exposed to the aquatic environment, and is susceptible to changes in water chemistry, including reduced oxygen levels (Simonis et al., 2024; Williams et al., 2019). Furthermore, OE’s neuroepithelium lacks vascularization, meaning that OSNs rely on passive oxygen diffusion from the environment to meet their high energetic demands (Bigdai et al., 2019). While the effects of hypoxia on brain function have been widely studied (Braga et al., 2013; Das et al., 2019; Park et al., 2021; Sawahata et al., 2021; Yu & Li, 2011), its impact on the olfactory system specifically remains largely unexplored. To our knowledge, only one recent study has directly examined the effects of hypoxia on olfactory function in fish. This report demonstrated reduced olfactory sensitivity to amino acids by electrophysiology (Tigert et al., 2025), but no study so far has explored the cellular or morphological basis of this dysfunction.

In this study, we aimed to investigate the functional and structural consequences of acute hypoxic insult to the olfactory system of adult zebrafish throughout the injury and recovery phases. For this, we conducted immunohistochemical, histological, biochemical and behavioral assays. We show that acute hypoxic exposure results in olfactory dysfunction associated with OE neurodegeneration and OB stress, followed by structural and functional recovery by 5 days. These findings further our understanding on the effects of low-oxygen conditions on the peripheral and central nervous system of a highly regenerative adult vertebrate.

## RESULTS

### Acute hypoxic exposure transiently impairs olfactory function

We sought to investigate the effects of acute hypoxic exposure on olfactory function and morphology in adult zebrafish. For this, fish were subjected to 15 min of severe hypoxia: 0.8 mg/L dissolved oxygen (DO) in a hypoxic chamber (Fig. 1A) and allowed to recover for 1 day post-hypoxia (dph) or 5 dph to assess recovery. Control fish were maintained in normoxic water (5-7 mg/l DO). Importantly, we did not observe any fish mortality using this exposure protocol. First, we assessed whether this hypoxic paradigm led to compromised olfactory function by analyzing olfactory-mediated responses to cadaverine, a colorless diamine released by decaying flesh that elicits robust and stereotypical aversive responses in zebrafish (Hussain et al., 2013). Single fish were acclimated to a rectangular tank and exposed to cadaverine for testing (Fig. 1B). We recorded swimming behaviors 30 seconds before and 30 seconds after odorant exposure and quantified three aversive behaviors: darting, freezing, and odor-evoked avoidance. To calculate the latter, we measured the cumulative time spent in the odor and non-odor halves of the arena. Importantly, the cadaverine solution remains in the odor half of the arena for the duration of the trial (Fig. 1C, highlighted in pink in Fig. 1D). Representative swimming trajectories are shown in Fig. 1D. Control fish displayed expected aversive behavioral responses to cadaverine, including darting (Fig. 1E), freezing (Fig. 1F), and avoidance (i.e., swimming to the non-odor half of the arena; Figs. 1G, D) (Vorhees et al., 2025).

**Figure 1.**
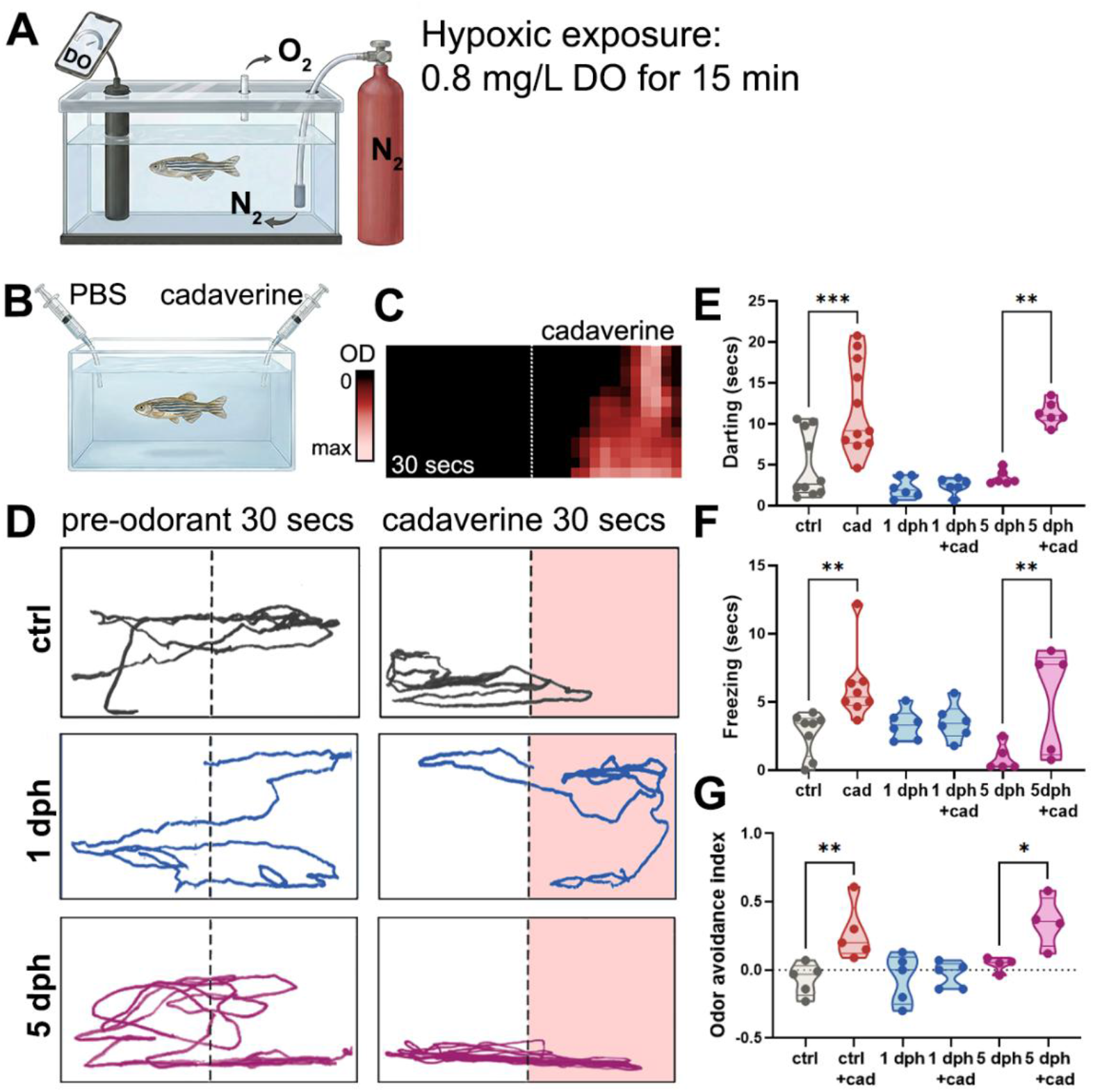
Odor-mediated behavioral responses to cadaverine in control and hypoxic fish. **(A)** Hypoxic exposure setup. We used a sealed hypoxic chamber with two ports. Nitrogen gas (N2) was delivered through a submerged inlet connected to an N2 cylinder into 300 ml of fish water. A second outlet tube allowed for excess gas exchange. DO levels were continuously monitored with a submerged digital DO probe. Once DO concentration reached 0.8 mg/L, nitrogen perfusion was stopped, and individual fish were placed in the sealed tank for 15 minutes. **(B)** Schematic of behavioral assay tank. PBS (control) and cadaverine (odorant) were dispensed into opposite sides of the tank. **(C)** Heatmap of cadaverine dispersal 30 seconds after delivery, indicating that cadaverine remains on one side of the behavioral tank. **(D)** Representative swimming trajectories of zebrafish 30 seconds pre-(left panels) and 30 seconds post-cadaverine administration (right panels, shaded in pink) in controls (top panels), 1 dph (middle panel), and 5 dph (bottom panels) fish (n= 5-8). **(E)** Time spent darting after cadaverine exposure in normoxic controls, 1 dph and 5 dph fish. ctrl vs. cad p = 0.0001; 5 dph vs. 5 dph cad p = 0.0012 (n= 6-10). **(F)** Quantification of time spent freezing after cadaverine exposure in 1 dph and 5 dph fish. ctrl vs. cad p = 0.0081; 5 dph vs. 5dph cad p = 0.0074 (n= 5-8). **(G)** Quantification of odor avoidance index in controls, 1 dph, and 5 dph fish. ctrl vs. cad p = 0.0058; 5 dph vs. 5dph cad p = 0.0244 (n= 4-5). Violin

In the 1 day post-hypoxia (1 dph) group, all these olfactory-mediated aversive responses to cadaverine were impaired (Figs. 1D-G), indicating reduced olfactory responses to cadaverine. Remarkably, by 5 days, fish regained olfactory sensitivity to cadaverine, demonstrated by restored darting, freezing, and avoidance behaviors (Figs. 1D-G). These results indicate that acute hypoxic exposure transiently impairs cadaverine-evoked aversive olfactory behaviors, with responses recovering by 5 days.

### Gross locomotion and spatial exploration are not affected by hypoxic treatment

To ensure that the effects on odor-mediated behavior were not confounded by alterations in general locomotor activity or exploratory activity caused by hypoxia, we quantified exploratory behaviors and gross locomotion parameters in control and hypoxic fish. First, we placed individual control and 1 dph fish into the same rectangular testing arena used for odorant assays and recorded swimming behaviors for 30 seconds without odorant delivery. We then analyzed exploratory behaviors using the recordings. For this, we sectioned the tank into thirds and quadrants to assess vertical and whole-tank exploration, respectively (Figs 2A, B). We quantified the cumulative vertical distance that fish swam, and the number of times the fish crossed into an adjacent quadrant. In both cases, we found no significant differences between groups (Figs. 2C, D), suggesting that exploratory behaviors in the experimental arena are largely unaffected by hypoxia exposure.

**Figure 2.**
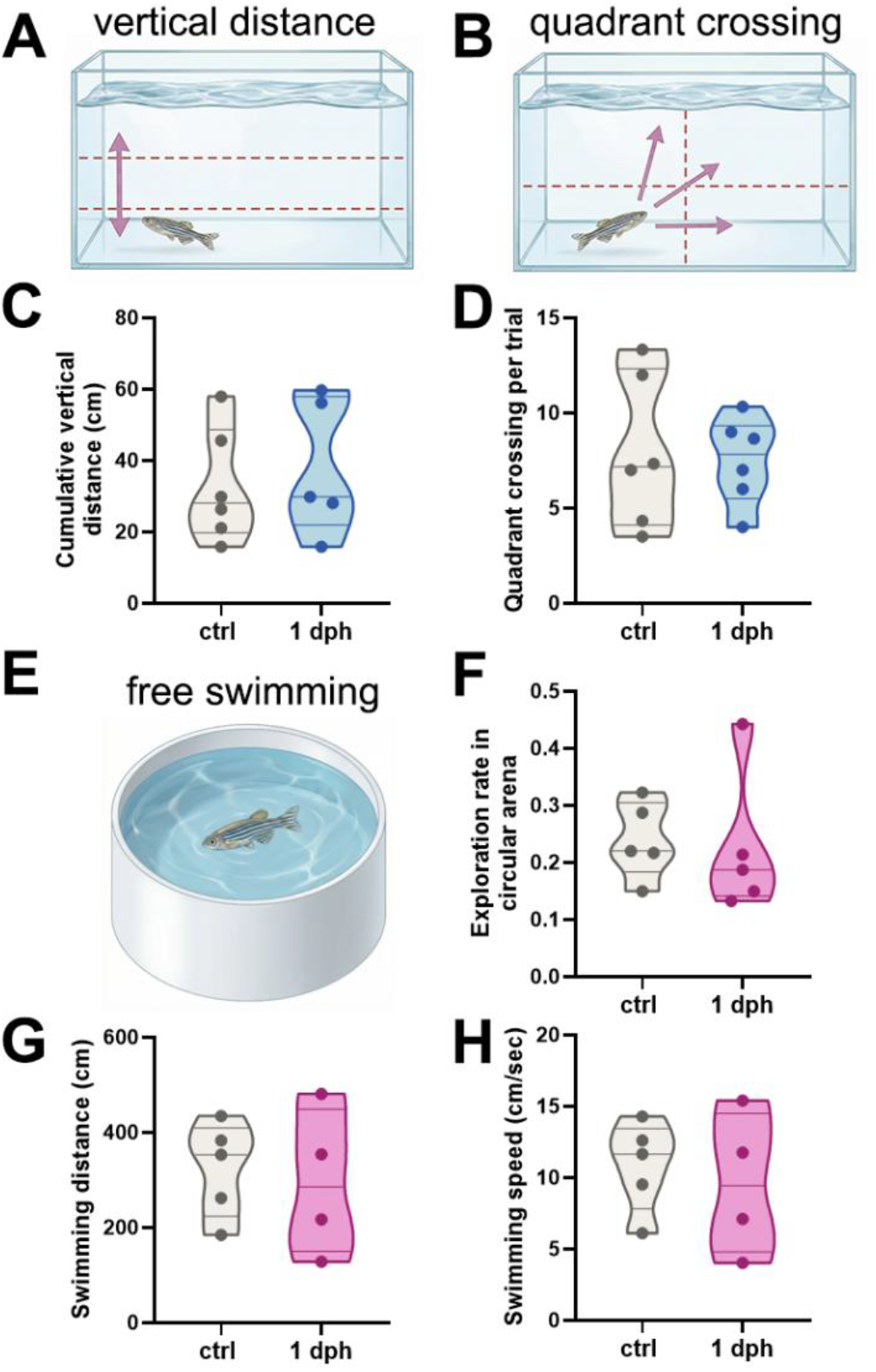
Exploratory behaviors and swimming parameters in control and hypoxic fish. **(A)** Diagram of the behavioral tank depicting three vertical zones used to quantify vertical distance displacement. **(B)** Behavioral tank schematic depicting four zones used to quantify zebrafish quadrant boundary crossings. **(C)** Quantification of cumulative vertical distance in controls and 1 dph fish (n= 6). **(D)** Quantification of quadrant crossing per trial in controls and 1 dph fish (n= 6). **(E)** Schematic of the behavioral circular arena used for free swimming assessment. **(F)** Quantification of exploration in a circular arena in controls and 1 dph fish (n= 5). **(G)** Quantification of swimming distance in a circular arena in controls and 1 dph fish (n= 5). **(H)** Quantification of swimming speed in a circular arena in controls and 1 dph fish (n= 5). Violin plots indicate mean, quartiles, and range.

Furthermore, we assessed whether hypoxia impairs gross locomotion. For this, individual fish were allowed to free-swim in a circular, larger arena (Fig. 2E). We calculated the exploration rate, and quantified the swimming speed and distance, and found no significant differences across groups (Figs. 2F-H). Collectively, these results indicate that the decreased responses to cadaverine observed are not attributable to locomotion or exploratory behavior impairments caused by hypoxia, but rather reflect deficits in odorant processing.

### Acute hypoxia causes transient neurodegeneration of the olfactory epithelium and alterations in nasal mucus

We predicted that hypoxic exposure is associated with structural alterations in the peripheral olfactory organ, and that these changes underlie the transient olfactory dysfunction observed. First, we examined the structural integrity of the sensory region of the OE, which is composed primarily of olfactory sensory neurons (OSNs), using immunohistochemistry (IHC) against HuC/D, a pan-neuronal marker. Our results revealed striking changes in the OE of hypoxic fish. We observed a significant thinning of the olfactory lamellae (Figs. 3A, B, C) along with a noticeable disruption of the HuC/D+ sensory layer of the epithelium (Fig. 3B), indicative of OSN loss. Echoing the recovery of olfactory-mediated behaviors by 5 dph, OSNs in the sensory epithelium regained their density and control morphology at this timepoint (Figs. 3A, B, C).

**Figure 3.**
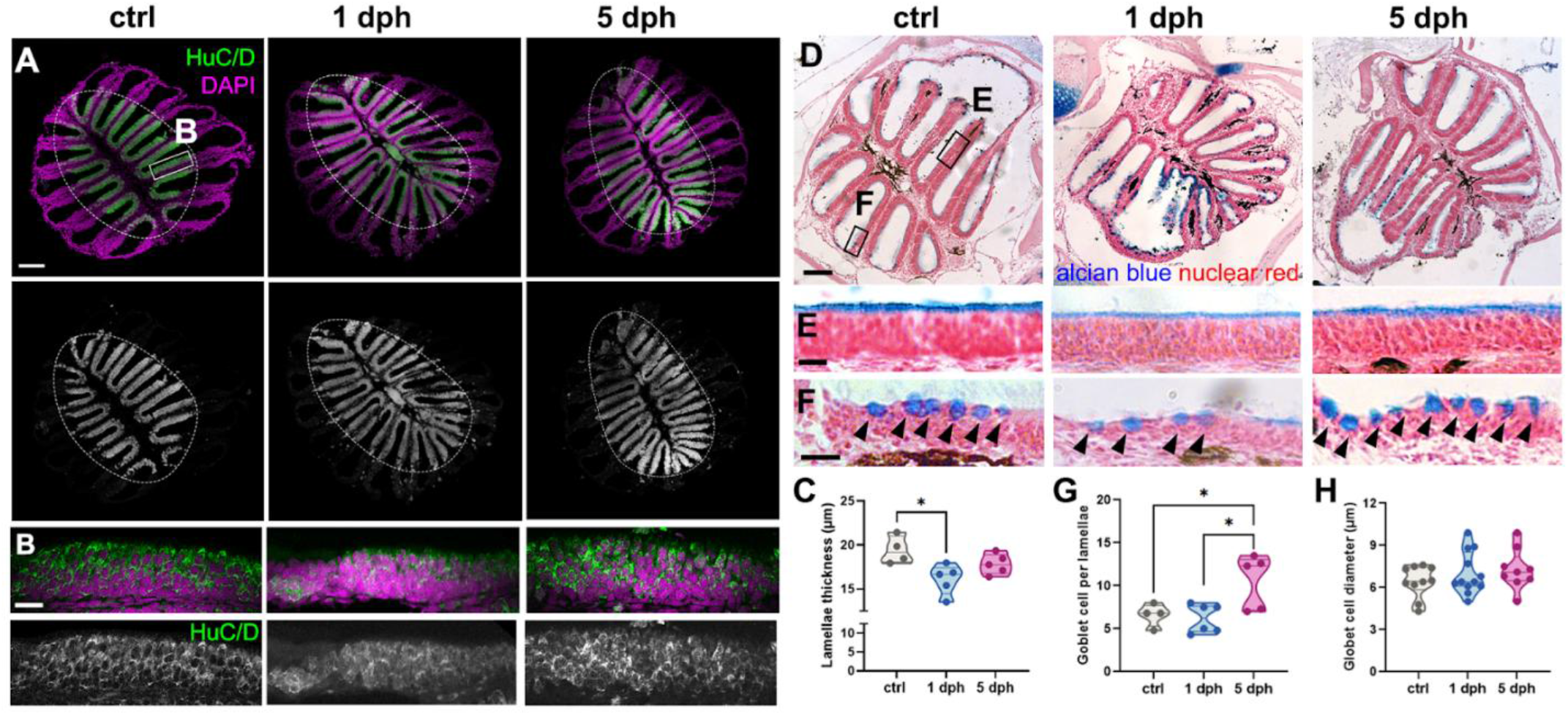
Effects of acute hypoxic exposure on OE structural integrity and the olfactory mucus layer. **(A)** Immunohistochemical stainings of HuC/D in OE sections from controls, 1 dph, and 5 dph fish. **(B)** Magnified views of the sensory epithelium (HuC/D+) of (A). **(C**) Quantification of olfactory lamellae thickness in controls, 1 dph and 5 dph groups (n= 4-5). ctrl vs. 1 dph p = 0.0162 9n= 4-5). **(D)** Alcian blue and nuclear fast red histological stainings of OE sections of controls, 1 dph and 5 dph fish. **(E)** Magnified views of the sensory epithelium of (D). The epithelial mucus layer is evident in blue staining. **(F)** Magnified views of the non-sensory epithelium of (D). Goblet cells are indicated by black arrowheads. **(G)** Number of goblet cells per olfactory lamellae in controls, 1 dph and 5 dph groups. ctrl vs. 5 dph. p = 0.0498; 1 dph vs 5 dph p = 0.0170 (n= 4 -6). **(H)** Quantification of goblet cell diameter (in μm) of controls, 1 dph and 5 dph fish (n= 9-12). Green: HuC/D; Magenta: DAPI. Scale bar: 100 μm in (A), 20 μm in (B), (C), and (D). Violin plots indicate mean, quartiles, and range. *p < 0.05.

Then, we examined the histological appearance of the epithelial mucus layer, which is important for olfactory signal transduction, as odorants must first dissolve in the mucus before binding to olfactory receptors on OSNs. For this, we performed histological stainings using Alcian blue and counterstained with fast nuclear red; the former stains acidic mucopolysaccharides and thus allows for the identification of mucins and mucus-producing cells. Nasal mucus alterations were assessed qualitatively, as tissue fixation and histological processing result in alterations of the mucus layer that preclude reliable quantification. We found that the mucus layer within the sensory epithelium appeared thinner and disorganized in 1 dph fish when compared to controls (Figs. 3D, E). In these magnifications of the histological staining, the significant thinning of the neuroepithelium (Figs. 1B, C) is also evident. Moreover, by 5 days, the mucus layer had returned to control appearance along with recovery of the epithelium (Figs. 3D, E). Furthermore, we examined mucus-producing goblet cells, identifiable by their characteristic rounded shape and strong Alcian blue positivity (Figs. 3D, F). Interestingly, although we did not observe changes in goblet cell numbers in 1 dph fish, we found elevated numbers at 5 dph compared to the other groups (Figs. 3F, G). Goblet cell diameter remained unchanged across experimental groups (Figs. 3F, G).

Together, these data indicate that acute hypoxia induces concurrent neurodegeneration of OSNs in the sensory epithelium and altered epithelial mucosal organization. The recovery of neuroepithelial thickness, along with the increase in the goblet cell population suggests compensatory processes in the olfactory organ as a result of the structural alterations caused by hypoxia.

### Peripheral leukocytic infiltration to the olfactory epithelium follows hypoxic exposure

Given that hypoxic exposure causes dramatic morphological changes and degeneration in the OE, we aimed to answer the question of whether this was related to inflammatory processes.

To study this, we examined the leukocytic response in the olfactory organ by means of an IHC against Lymphocyte Cytosolic Protein 1 (Lcp1), a pan-leukocytic marker. Our group and others have reported that Lcp1+ cells, primarily neutrophils, are constitutively present in the outer epithelial folds of the olfactory organ (Figs. 4A, B) (Palominos et al., 2022). It is not possible to count individual leukocytes in this region due to their high abundance; thus, we quantified the optical density of Lcp1 signal within the entire olfactory lamellae to assess leukocyte density. Using this method, we found a significant increase in Lcp1 signal in 1 dph fish when compared to controls, with signal returning to control levels in 5 dph groups (Figs. 4A, B, C, D). Upon further examination, we noticed an increase in the number of Lcp1+ cells in the sensory lamellae. To confirm this, we performed a double IHC staining against HuC/D and Lcp1 and found an increased number of individual Lcp1+ cells found within the HuC/D+ sensory epithelium (Figs. 4D, E). Notably, in control fish, most leukocytes are found underlying the basal lamina propria of the epithelium (Fig. 4D, indicated by asterisks), while in the 1 dph group, an increased number of Lcp1+ cells were observed infiltrating apical layers, surrounding OSNs (Fig. 4D, indicated by white arrowheads). We also noticed changes in leukocyte morphology in the 1 dph group: resting leukocytes in control fish were small and rounded, whereas activated leukocytes in hypoxic fish adopted an enlarged and/or ameboid phenotype (Figs. 4B, D), consistent with cytoplasmic extensions that facilitate migration and tissue infiltration previously reported (Chen et al., 2022; Palominos et al., 2022). Conversely, the inflammatory response subsided within 5 days, as indicated by Lcp1+ cell density and presence in the sensory lamellae returning to control levels in the 5 dph group (Figs. 4B, C, D, E). Collectively, these results are indicative of increased inflammatory processes caused by hypoxia, as demonstrated by leukocytic recruitment throughout the entirety of the olfactory organ, including the OE’s OSN-rich sensory region.

**Figure 4.**
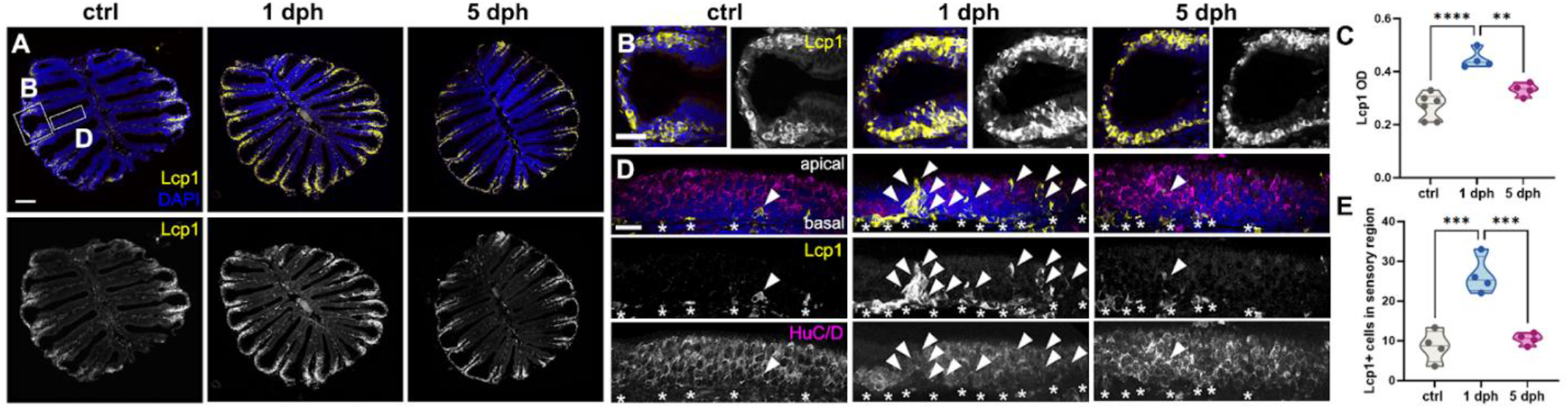
Effects of hypoxia on leukocytic recruiting to the OE. **(A)** Immunohistochemical staining of Lcp1 in OE sections from controls, 1 dph, and 5 dph fish. (**B)** Magnified views of the outer non-sensory epithelial fold of (A). **(C)** Quantification of Lcp1 optical density (OD) in sections from (A). ctrl vs. 1 dph p <0.0001; 1 dph vs. 5 dph p = 0.0051 (n= 4-6).. **(D)** Magnified views of the sensory epithelium of OE sections double immunohistochemically stained against HuC/D and Lcp1. Lcp1+ cells present in the basal membrane are indicated by asterisks, while white arrowheads indicate Lcp1+ leukocytes infiltrating apical sensory layers. **(E)** Quantification of Lcp1+ cells found in the sensory lamellae of ctrl, 1 dph, and 5 dph groups. ctrl vs. 1 dph p = 0.0002; 1 dph vs. 5 dph p = 0.0005 (n = 4). Yellow: Lcp1; Magenta: HuC/D. Scale bar: 100 μm in (A), 20 μm in (B), and 40 μm in (D). Violin plots indicate mean, quartiles, and range.**p < 0.01, ***p < 0.001, ****p < 0.0001.

### Acute hypoxia induces central reactive astrogliosis and mitochondrial dysfunction in the olfactory bulb

Having established that acute hypoxia is associated with structural dysregulation and inflammation of the OE, we next sought to identify inflammatory and metabolic changes in the olfactory bulb (OB) that may be related to OE changes. First, we investigated reactive astrogliosis by means of an IHC against Glial Fibrillary Acidic Protein (GFAP), a marker of astroglia and olfactory ensheathing cells (OECs, a specialized glial population associated with the olfactory nerve layer (ONL) in the OB (Lazzari et al., 2014). Our results revealed a significant increase in GFAP immunoreactivity in the OB at 1 dph compared to normoxic controls, particularly along the olfactory nerve layer (ONL; Fig. 5A), indicative of robust astroglial activation following hypoxic exposure (Figs. 5A, B). By 5 dph, GFAP immunoreactivity returned to control levels, suggesting resolution of the astroglial response in parallel with behavioral recovery (Figs. 5A, B).

**Figure 5.**
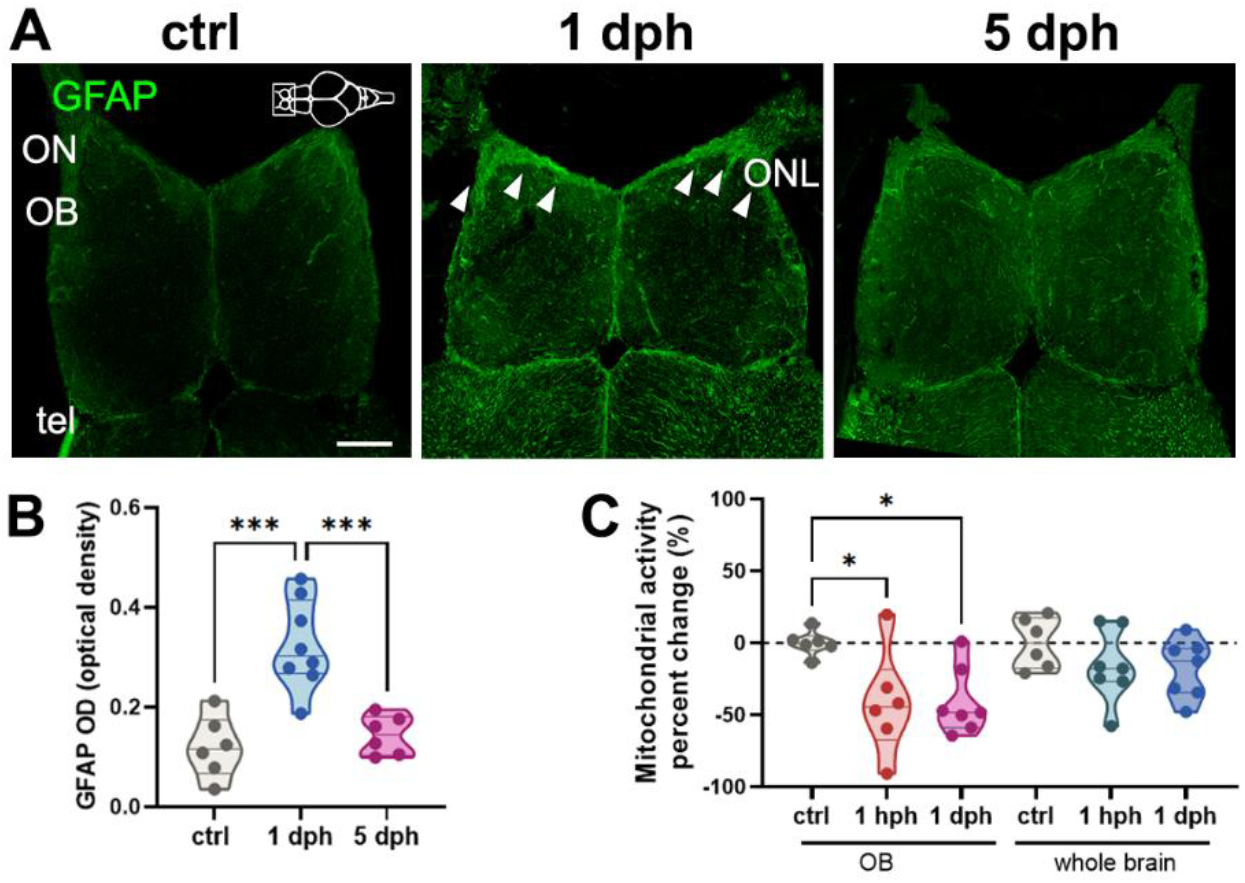
Effects of hypoxia on reactive astrogliosis and mitochondrial activity in the OB. **(A)** Glial Fibrillary Acidic Protein (GFAP) immunohistochemical stainings of horizontal sections of the OB from controls, 1 dph, and 5 dph fish. Increased GFAP staining along the olfactory nerve and the olfactory nerve layer in the OB is indicated with white arrowheads. **(B)** Quantification of GFAP optical density (OD) in OB sections from (A). ctrl vs. 1 dph p = 0.0001; 1 dph vs. 5 dph p = 0.0005 (n=6-8). **(C)** Quantification of mitochondrial activity percent change in the OB and whole brains of ctrl, 1 hph, and 1 dph groups. OB: ctrl vs. 1 hph p = 0.0442; ctrl vs. 1 dph p = 0.0393 (n = 6-7). Green: GFAP. Scale bar, 100 μm. Violin plots indicate mean, quartiles, and range. *p < 0.05, ***p < 0.001.

Exposure to hypoxia can compromise mitochondrial function and integrity due to increased oxidative stress, reduced ATP production, and altered cellular metabolism. Thus, we asked whether acute exposure compromises mitochondrial function in the OB. We employed a colorimetric assay widely used as an index of mitochondrial dehydrogenase activity in intact brain sections of zebrafish (Yu & Li, 2011). This assay indicates mitochondrial activity through the reduction of a colorless 2,3,5-triphenyltetrazolium chloride (TTC) solution to red triphenyl formazan. Given that a reduction of mitochondrial activity could be rapid and transient, we quantified TTC staining after 1 hour (1 hph) and 1 day post hypoxic exposure (1 dph) by measuring the optical density (OD) of the OB and the rest of the brain. We then converted these values to percent change in mitochondrial activity. OB from 1 hph and 1 dph fish displayed larger unstained areas and significantly higher luminosity values compared to controls, indicating a marked reduction in mitochondrial activity following hypoxic exposure (Fig. 5C). Interestingly, mitochondrial activity levels of the rest of the brain of hypoxia-exposed fish did not differ from controls (Fig. 5C). These findings suggest that the OB is particularly susceptible to low-oxygen conditions, leading to compromised mitochondrial function early after exposure. Our data demonstrate that acute hypoxia induces coincident reactive astrogliosis and mitochondrial dysfunction in the OB.

### Hypoxic exposure induces cell proliferation within the OE and telencephalic ventricular zone

Unlike many vertebrates, zebrafish retain extensive neuroregenerative capacity throughout adulthood, enabling heightened cell proliferation and increased resilience after neural injury. This ability is driven by progenitor cell populations’ capacity to rapidly respond to injury and promote recovery. Hence, we hypothesized that the post-hypoxia functional and morphological recovery described herein is sustained by cell proliferation. To study this, we used Proliferating Cellular Nuclear Antigen (PCNA) immunostainings to visualize actively proliferating cells within the OE and the ventral the telencephalon, a well-described neurogenic niche posterior to the OB (Adolf et al., 2006). Specifically, we looked at two proliferative zones within the VZ, the ventroventral (Vv) and ventrodorsal (Vd) regions, which serve as a critical neurogenic niche driving regeneration and repair of the OB following injury (Diotel et al., 2015; Marz et al., 2010; März et al., 2011).

Control fish exhibited constitutive cell proliferation in the OE, in particular in the outer epithelial fold (Figs. 6A, B). Our results indicate that hypoxia induces cell proliferation within the olfactory epithelium, as evidenced by a significant increase in PCNA+ cells at 1 dph when compared to controls (Fig. 6A, B, C). Interestingly, PCNA immunoreactivity increased within the sensory region and the interlamellar curve (ILC) in the 1 dph group (Fig. 6B, indicated by dashed lines). Moreover, PCNA+ numbers returned to control levels by 5 dph (Figs. 6A, B, C). Similarly, we noticed a significant increase in cell proliferation within the ventral telencephalon (Vv) of 1 dph fish when compared to controls (Fig. 6D, E). However, we found no difference in PCNA+ profiles in the Vv between the two hypoxic groups (Figs. 6D, E), indicating that proliferation in the ventral telencephalon remains sustained at 5 dph. Additionally, no differences were found in the ventrodorsal region (Vd) across groups (Fig. 6D, F).

**Figure 6.**
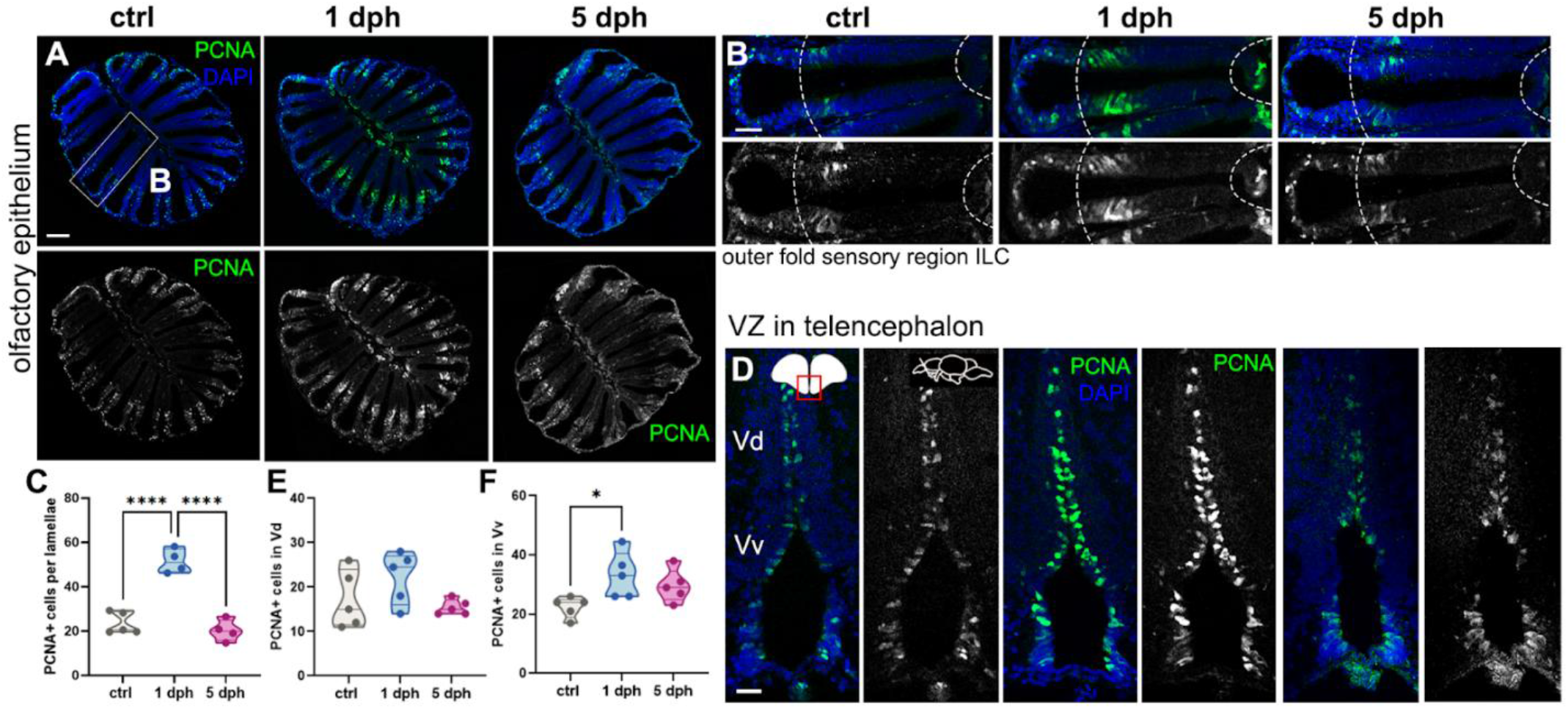
Effects of acute hypoxic exposure on cell proliferation in the OE and telencephalon. **(A)** Proliferating Cellular Nuclear Antigen (PCNA) immunohistochemical stainings of OE sections from controls, 1 dph, and 5 dph fish. **(B)** Magnified views from (A). Dashed lines indicate the following anatomical divisions of the lamellae: outer non-sensory epithelial fold (left), sensory region (middle) and intralamellar curve (ILC, right). **(C)** Quantification of PCNA+ cells in single olfactory lamellae from (A). ctrl vs. 1 dph p < 0.0001; 1 dph vs. 5 dph p < 0.0001(n= 4-5). **(D)** PCNA immunohistochemical stainings of coronal telencephalon sections from controls, 1 dph, and 5 dph groups. The ventrodorsal (Vd) and ventroventral (Vv) regions within the ventricular zone (VZ) are indicated. **(E)** Quantification of PCNA+ cells within the Vd telencephalic region shown in (**C)** (n= 5). **(F)** Quantification of PCNA+ cells within the Vv telencephalic region from (C). ctrl vs. 1 dph p = 0.0329 (n = 5). Violin plots indicate mean, quartiles, and range. Green: PCNA; Blue: DAPI. Scale bar, 100 μm in (A) and (D); 20 μm in (B). Violin plots indicate mean, quartiles, and range.*p < 0.05, ****p < 0.0001.

Collectively, these results suggest that the structural and functional recovery of olfactory function observed at 5 dph could be associated with rapid cellular proliferation in the OE. Furthermore, the differences in proliferation rates observed across telencephalic regions and recovery times suggest a complex post-hypoxic response by distinct cell populations within the VZ.

## DISCUSSION

In this study, we investigated the functional and structural effects of acute oxygen deprivation on the olfactory system. Adult zebrafish is an advantageous model organism for studying hypoxia-induced repair responses because they are able to withstand low-oxygen conditions and exhibit robust widespread regeneration. Our work reveals that acute hypoxic insult transiently impairs olfactory function through concurrent degenerative processes in the OE and metabolic dysfunction in the OB, followed by rapid recovery mediated by compensatory cell proliferation across the olfactory axis.

### Acute hypoxia transiently impairs olfactory-mediated behavior without affecting gross locomotion

We show that acute hypoxia causes olfactory dysfunction, as demonstrated by reduced responses to cadaverine at 1-day post hypoxia (dph). Cadaverine is produced during tissue decomposition and activates TAAR13c-expressing OSNs, eliciting strong odorant-evoked aversive responses (Hussain et al., 2013). Notably, olfactory-mediated responses to cadaverine were fully restored at 5 dph, indicating that olfactory dysfunction is reversible.

Importantly, our hypoxic paradigm does not impair swimming or exploratory behaviors, indicating that the impaired responses to cadaverine arise due to olfactory loss rather than locomotor impairment. This is consistent with prior reports showing that, while motor deficits are observed at very early recovery timepoints (between 1 and 2 hph) locomotion normalizes as soon as 3 hph (Braga et al., 2013; Yunkyoung Lee et al., 2018; Marino et al., 2020). This temporal pattern suggests that hypoxia-mediated motor deficits may reflect transient neuromuscular metabolic stress rather than impairment of neuronal motor circuits (Cadiz et al., 2019). Normal swimming resume as compensatory responses such as improved oxygen usage are deployed (Esbaugh et al., 2009; Van Ginneken et al., 1995).

Our findings align with reports of post-hypoxia olfactory decline across different species (Bigdaj et al., 2018; Kim et al., 2023). Mice exposed to chronic intermittent hypoxia exhibited broad olfactory alterations (Hernandez-Soto et al., 2021). Similarly, humans exposed to hypoxic conditions due to sleep apnea or experimental high altitude, experience olfactory dysfunction, as evidenced by a lowered olfactory threshold (Binar & Gokgoz, 2021; Huppertz et al., 2018). Our study extends these findings by providing a characterization of post-hypoxia cellular and morphological changes across the olfactory axis.

Importantly, there are some discrepancies in the literature regarding adult zebrafish mortality after hypoxic exposure. We did not observe any fish mortality with our hypoxic exposure paradigm of 0.8 mg/l DO for 15 minutes, consistent with studies reporting survival under similar conditions from 10 minutes for up to 48 hours (Marino et al., 2020; Rees et al., 2001; Roesner et al., 2006). However, other reports have described elevated mortality under similar parameters (Yunkyoung Lee et al., 2018; Yu & Li, 2011). These discrepancies may be explained by the use of different zebrafish strains, methodological differences, and/or water chemistry. Our paradigm was carefully validated to produce a severe but sublethal acute hypoxic insult.

This work focused on studying olfactory-evoked responses to cadaverine, but whether hypoxia impairs olfactory responses to other odorant categories remains to be explored. Notably, moderate hypoxia reduces olfactory sensitivity to amino acids in the teleost gilt-head seabream (*Sparus aurata*) (Tigert et al., 2025), suggesting that hypoxia could lead to reduced food-seeking behaviors. Exploring how hypoxia affects a wider range of olfactory-evoked behaviors, including foraging and social interactions, can further our understanding of the broader deleterious impact of low-oxygen conditions in aquatic animals. The cellular and morphological mechanisms underlying this transient olfactory dysfunction are discussed in the following sections.

### Peripheral neurodegeneration and mucus disruption impair olfactory function

The structural integrity of the olfactory organ is crucial for olfactory signal transduction. Our findings revealed a significant thinning of the sensory OE lamellae at 1 dph, indicative of OSN loss and consistent with the deficits in olfactory function observed at this timepoint. These structural alterations recovered by 5 dph, paralleling the restoration of olfactory-mediated behaviors. Our findings confirm results from a study in mice that described a post-hypoxia reduced expression of olfactory marker genes in the OE (Kim et al., 2023).

The OE is particularly susceptible to low oxygen because it is a neuron-rich tissue that relies heavily on diffusion from the external environment rather than a direct blood supply (Bigdai et al., 2019). Olfactory signal transduction is an energetically demanding process that relies on robust ATP production in the dendritic knob and processing of extracellular glucose in odorant cilia (Villar et al., 2017). Thus, a reduction in available oxygen in OSNs could be expected to impair oxidative phosphorylation and reduce ATP production, thereby diminishing olfactory sensitivity. This is consistent with a report showing that hypoxia reduces amino acid-evoked OSN potentials suggesting that the initial stages of odorant recognition are impaired by low oxygen, possibly through ATP reduction (Tigert et al., 2025). A limitation of our study is that we did not assess mitochondrial activity in the OE, because olfactory organs pigment can confound the results of colorimetric TTC assays. In future experiments, other biochemical assays could be used to assess mitochondrial function in OSNs.

In addition to OSN degeneration, we propose that the disruption of the nasal mucus layer might a contributing factor on the olfactory dysfunction observed. The mucus layer serves two important functions: it acts as a medium for the diffusion of both odorants and oxygen. Odorants must dissolve in nasal mucus before accessing olfactory receptors within OSN cilia (Rygg et al., 2013), while dissolved oxygen from the water diffuses to the mucus layer to the underlying neuroepithelium (Cox, 2008), thus the thickness and composition of nasal mucus directly affects the rate of diffusion of these important solutes. Moreover, sustentacular cells in the OE release glucose to the mucus layer where olfactory cilia incorporate it as energy source (Nunez-Parra et al., 2011). Thus, disruption of the nasal mucus could further impair OSN metabolic activity. Therefore, we propose that hypoxia-induced disruption of the mucus layer may both impair odorant transduction and further exacerbate the oxygen deficit experienced by OSNs.

While we were unable to determine the thickness of the mucus layer with our current techniques, our qualitative examination revealed histological changes within this layer as supported by our findings on goblet cell number. Interestingly, we observed an increase in the number of goblet cells at a timepoint when the OE’s morphology had already recovered. This delayed goblet cell expansion suggests that mucosal remodeling continues beyond the window of hypoxic insult, possibly reflecting a compensatory response to the epithelial damage. Notably, while research on the impact of low oxygen in olfactory mucus-producing glands is scarce, prolonged hypoxia has been associated with increased mucus production and rearrangement of mucus-secreting cells in other epithelial tissues (Polosukhin et al., 2011), consistent with our findings.

Given the scarcity of reports regarding the nasal mucus dynamics in the teleost olfactory organ, our findings are an important contribution to this underexplored area of olfactory physiology.

### Transient inflammation in the OE coincides with functional and structural recovery

The robust leukocytic response in the peripheral olfactory organ following hypoxic exposure reflects the dynamic role that immune cells play in response to injury and supporting neural regeneration in the peripheral nervous system (PNS) in zebrafish (Villegas et al., 2012), including the olfactory organ (Palominos et al., 2022).

Neutrophils are the most abundant type of circulating leukocytes and are typically the first immune cells recruited to sites of tissue injury (Harvie & Huttenlocher, 2015; Mokhtar et al., 2023). The olfactory organ of zebrafish contains a resident population of neutrophils localized in the outer external folds and within the basal lamina (Palominos et al., 2022). Importantly, macrophages are also recruited following PSN injury to aid in debris removal and repair (Calvo-Ochoa et al., 2025; Carrillo et al., 2016; Var & Byrd-Jacobs, 2020).

The increase in Lcp1+ density following hypoxia, the infiltration into the neuroepithelium, and changes in cell morphology indicate leukocytic recruitment, consistent with prior reports of neutrophils and macrophages migration to the injured OE (Calvo-Ochoa et al., 2025; Palominos et al., 2022). A potential mechanism by which hypoxia might promote leukocytic migration to the OE is through the hypoxia-inducible factor (HIF) signaling. This pathway leads to upregulation of genes that promote leukocyte recruitment, survival and immune activity (He et al., 2022; Walmsley et al., 2005). Future work could explore whether this signaling pathway is responsible for the immune response observed herein.

While inflammation is essential for initiating tissue repair, its resolution is required to enable regenerative processes. Our findings show that the inflammatory response subsides by 5 days, concurrent with OE’s morphological recovery. The transient nature of the leukocytic response is consistent with evidence of acute inflammation as a key regulator of neuronal repair and regeneration in zebrafish (Var & Byrd-Jacobs, 2020). Acute inflammation has been shown to enhance proliferation of neural progenitors and subsequent neurogenesis following injury in different regions of the zebrafish brain, spinal cord, and PNS (Carrillo et al., 2016; Kyritsis et al., 2012; Tsarouchas et al., 2018; Villegas et al., 2012). On the other hand, our understanding of the role of leukocytes in the OE remains limited. In mice, OE-infiltrating neutrophils express neurogenesis-related genes following damage (Ogawa et al., 2021), while attenuation of inflammatory processes in the injured OE impaired cell proliferation (Chen et al., 2017). This raises the possibility that the leukocytic response observed might support repair and regeneration of lost OSNs. Future research studying whether modulating the immune response impairs regeneration in the olfactory system is needed to test this hypothesis.

While leukocytic infiltration reflects the peripheral inflammatory response to hypoxia, concurrent changes in the OB suggest that the injury extends beyond the epithelium, affecting central olfactory processing, as described below.

### Early mitochondrial dysfunction and astrogliosis reveal inherent susceptibility of the OB

Our findings show that post-hypoxia olfactory dysfunction is associated with structural and metabolic alterations along the olfactory axis. Our TTC data reveal that mitochondrial dehydrogenase activity in the OB is reduced as early as 1 hour post hypoxia and that it continues to be reduced throughout the first day, while mitochondrial activity in the rest of the brain remains unchanged. This suggests that the OB has an inherent susceptibility to hypoxic insult. Other reports have shown that that severe hypoxic exposure decreases mitochondrial dehydrogenase activity in the telencephalon and optic tectum, with mitochondrial activity remaining reduced for up to 24 hours before normalizing by 48 hours post-hypoxia (Braga et al., 2013; Braga et al., 2016). Furthermore, repeated hypoxic exposure triggers downregulation of brain glucose levels and several glucose metabolites, with supplementation of glucose and glucosamine restoring learning and memory impairment (Park et al., 2021). Additionally, hypoxia changes the acetylome of brain proteins, indicating changes in redox status and changes in electron transport activity (Dhillon & Richards, 2018). Collectively, these results demonstrate that low oxygen widely impairs oxidative phosphorylation and glycolytic processes in the zebrafish brain. Our work extends these findings by demonstrating that the OB exhibits inherent vulnerability to low oxygen, pointing to differences in metabolic demands or redox protective mechanisms across brain regions.

Hypoxia suppresses mitochondrial oxidative phosphorylation, reducing ATP production and increasing AMP levels, which collectively impair energy-demanding neuronal processes (Allen et al., 2020; Dhillon & Richards, 2018; Douglas et al., 2010; Fuhrmann & Brüne, 2017). In the context of the OB, where odorant-evoked synaptic transmission in the glomerular layer is particularly oxygen-consuming (Lecoq et al., 2009; Nawroth et al., 2007), this metabolic disruption would be expected to impair olfactory signal processing, amplifying the peripheral deficit caused by OSN degeneration in the OE.

Concurrent with mitochondrial dysfunction, we observed a significant increase in GFAP immunoreactivity at 1 dph, particularly along the olfactory nerve layer (ONL), with resolution by 5 dph. Our results add to the existing literature showing that hypoxia causes reactive gliosis in the telencephalon (Yunkyoung Lee et al., 2018). GFAP is expressed by multiple glial populations in the OB, including astroglia and olfactory ensheathing cells (OECs), the latter being particularly concentrated in the ONL where we observed the greatest increase in signal at 1 dph (Lazzari et al., 2014). OECs are specialized glia that envelope OSN axons as they project from the OE to the OB and are known to support olfactory neurogenesis and axonal regeneration (Chuah & West, 2002; Franklin & Barnett, 1997; Raisman & Li, 2007). Therefore, the increase in GFAP immunoreactivity we observed along the ONL may reflect activation of OECs in response to OSN axonal degeneration in the OE, rather than, or in addition to, classical reactive astrogliosis. Future studies employing markers specific to OECs, such as S100β, would help disambiguate the relative contributions of astrocytes and OECs to the glial response observed following hypoxia.

Importantly, astroglia can exert both neuroprotective and neuroinflammatory effects in response to hypoxia, releasing neurotrophic factors and antioxidants that protect neurons from hypoxic injury (Marina et al., 2016), while also producing pro-inflammatory cytokines and reactive oxygen species that may exacerbate damage (Merelli et al., 2021). The resolution of GFAP immunoreactivity by 5 dph, coinciding with behavioral recovery, suggests that in our model the astroglial response is ultimately protective, consistent with other injury models in which astrogliosis has been reported as neuroprotective or regenerative (Baumgart et al., 2012; Kroehne et al., 2011b; Pellegrini et al., 2007; Skaggs et al., 2014).

The resolution of the post-hypoxic peripheral degeneration and central metabolic dysfunction reported appears to be driven by compensatory cell proliferation across the axis, as discussed in the following section.

### Compensatory cell proliferation underlies rapid recovery

The remarkable and rapid recovery of olfactory function by 5 dph – despite the severity of the peripheral and central disruption observed at 1 dph — points to extensive compensatory mechanisms actively driving structural repair across the olfactory axis.

The robust increase in cell proliferation throughout the OE supports the hypothesis that neurogenic mechanisms occur after hypoxic insult to replace damaged or lost OSNs. This damage-responsive proliferative response is consistent with the well-documented abilities of progenitor cell populations to drive neuronal replenishment (Calvo-Ochoa et al., 2025; Kocagöz et al., 2022; Sireci et al., 2024). Resolution of the proliferative response by 5 dph when the neuroepithelium is structurally and functionally restored, further indicates that this is a transient, injury-driven response.

Proliferating PCNA+ profiles in control fish were localized within the outer epithelial folds, where globose basal cells (GBCs) give rise to constitutive maintenance proliferation (Bayramli et al., 2017; Kocagöz et al., 2022). Following hypoxia, proliferating cells were found across the OE, including within the sensory region and the ILC, areas enriched with horizontal basal cells (HBCs), progenitors that contribute to damage-induced proliferation (Sireci et al., 2024). Future work will be needed to parse out which progenitors are activated by hypoxic exposure.

Concurrent with the peripheral proliferative response, we observed a robust increase in PCNA+ cells within the ventroventral (Vv) region of the rostral telencephalon. Interestingly, proliferation in this region remained increased at 5 dph. The Vv is an active neurogenic niche densely packed with fast-cycling progenitors that constitutively give rise to proliferating neuroblasts that migrate towards the OB through a rostral migratory stream, similar to the one described in mammals (Adolf et al., 2006; Grandel et al., 2006; Kishimoto et al., 2011).

In contrast to OE proliferation, which subsides rapidly by 5 dph, the sustained activation of the Vv has important implications for supporting OB recovery. We posit that this prolonged response reflects the need for longer-term remodeling of OB neurons needed to replace olfactory glomerular connectivity disrupted by the transient loss of OSN afferents. While constitutive and post-injury neuronal replacement in the OE occurs quickly (Iqbal & Byrd-Jacobs, 2010; Kocagöz et al., 2022), newborn bulbar neurons generated in the Vv take several days to weeks to reach the OB (Calvo-Ochoa et al., 2025; Godoy et al., 2020; Kishimoto et al., 2011; Trimpe & Byrd-Jacobs, 2016). Furthermore, there were no changes in cell proliferation within the ventrodorsal (Vd) region. This differential proliferation profile is consistent with the distinct progenitor identities of both proliferative niches: while the Vv is populated primarily by fast-cycling neuroblast progenitors, the Vd is composed mostly of slow-cycling progenitors whose output is directed toward telencephalic regions (Grandel et al., 2006; Kizil et al., 2012; Kroehne et al., 2011a; Rothenaigner et al., 2011). The selective activation of Vv progenitors suggests that our sublethal hypoxic insult model may be insufficient to recruit slow-cycling populations that might respond to more severe injury signals to undergo proliferation.

A study that used a comparable hypoxic exposure reported cell proliferation in the Vd domain. This discrepancy is likely explained by methodological differences: their hypoxic paradigm was considerably more severe, resulting in increased mortality; and proliferation was assessed by BrdU labeling and not PCNA (Yunkyoung Lee et al., 2018). These differences suggest that the extent of brain damage associated with varying degrees of hypoxia severity is a determinant in the selective engagement of fast-vs slow-cycling progenitors in the telencephalon.

Hypoxia regulates proliferation in the zebrafish nervous system via Wnt/β-catenin signaling (Demirci et al., 2020; Meyers et al., 2012; Shimizu et al., 2018; Shitasako et al., 2017), and most notably, in the damaged OE (Kocagöz et al., 2022). Future work could elucidate the role of Wnt/β-catenin signaling in the proliferative response to hypoxia.

Collectively, these findings indicate that the rapid recovery of olfactory function following hypoxic insult arises from a coordinated repair response that engages the proliferation of progenitor populations and the timely resolution of inflammatory signaling across the olfactory system.

## Conclusions

This study provides the first comprehensive characterization of the cellular and morphological mechanisms underlying hypoxia-induced olfactory dysfunction. We extend prior work by revealing the structural basis for post-hypoxic injury and repair across the olfactory axis in a model organism known for its remarkable regenerative capacity, the adult zebrafish. Further investigation of repair and regeneration processes described here may ultimately inform therapeutic strategies for enhancing neural repair following hypoxic-ischemic brain injury.

## ACKNOWLEDGEMENTS

We are grateful to current and past members of the Calvo lab for support throughout the completion of these studies; in particular to Lexus Putt, Ethan DeKoker, and Marco Lopez-Vargas. We are grateful to our late colleague Ron Fleischmann for meaningful discussions during the conceptualization stage of this research project. Research reported in this publication was supported in part by funding provided by the National Aeronautics and Space Administration (NASA), under award number 80NSSC25M7087, Michigan Space Grant Consortium (MSGC) through undergraduate research grants to SLD, KED, and EAT. ECO and LZ were supported by the National Science Foundation (grant BRC-BIO 2313303), and Hope College.

## Author contributions

ECO conceived the project and designed experiments. SLD, KED, and EAT performed hypoxic exposure, tissue preparation, sectioning, antibody stainings, confocal microscopy, and analyzed imaging data. CJH, CLL, LAH, KED, LS, and SLD performed behavioral assays and analyzed behavioral data. KED and LS performed histological stainings, light microscopy, and analyzed imaging data. ECO, SLD, and KED contributed to writing and editing the manuscript. ECO, SLD, KED, and EAT acquired funding.

## MATERIALS AND METHODS

### Animals

Adult wild-type zebrafish (*Danio rerio)* of both sexes were kept and bred in an aquatic system at 28º C (Aquaneering, San Diego, CA) located at Hope College’s zebrafish facility. The facility was kept on a 12-hour light/dark cycle. Routine feedings consisted of pellet fish food (Reef Nutrition, Campbell, CA) twice daily and fresh brine shrimp (Brine Shrimp Direct, Ogden UT) once daily. All experimental procedures were performed in accordance with the National Institutes of Health Guide for the Care and Use of Laboratory Animals (NIH Publication No. 80-23) and were approved by Hope College’s Institutional Animal Care and Use Committee (HCACUC). All efforts were made to minimize the number of animals used.

### Acute hypoxic exposure

Fish were exposed to hypoxic conditions in a hypoxic chamber formed by a sealed 7 in × 5 in × 3.5 in glass container (Rubbermaid, Atlanta, GA) outfitted with two gas ports. To displace dissolved oxygen (DO) in 300 ml of fish water, nitrogen gas (N_2_) was delivered through a submerged surgical tube connected to an N_2_ cylinder. A second outlet tube was positioned at the water surface to allow for excess gas exchange and maintain controlled circulation within the chamber. DO levels were continuously monitored with a submerged digital DO probe (RCYAGO, China). (Fig. 1) Once the DO concentration reached 0.8 mg/L, nitrogen perfusion was stopped (Yu & Li, 2011), and individual fish were placed in the sealed tank for 15 minutes. Following hypoxic exposure, fish were allowed to recover for one or five days post hypoxia (dph) before being used for experiments.

### Olfactory-mediated behavioral assays

We used rectangular clear tanks (4.7 cm × 9.6 cm × 15.8 cm) (AquaCulture, Fairfield, NJ) to assess behavioral responses to the odorant cadaverine. Each tank was filled with 1.5 liters of fresh water from the zebrafish system in our facility. The tanks were fitted with surgical plastic tubing placed on the opposite side of the tank, slightly below the water level to avoid water disruption. Syringes were attached to the tubes and used for the simultaneous administration of 1 ml of odorant and control (PBS) solutions on opposite sides of the tank. Tubes and syringes used for cadaverine delivery were not used for PBS exposure to avoid cross-contamination. We used a digital camera to record fish swimming behaviors, placed behind a white panel that surrounded the tank to conceal the presence of investigators, in order to minimize fish distraction during testing (Vorhees et al., 2025).

Before experiments, fish were fasted in isolation for 48 hours to heighten olfactory sensitivity. On the day of the experiment, individual fish were placed in the experimental tank and acclimated for an hour total, with 30 minutes in the absence of noise. Fish were recorded for 30 seconds before and 30 seconds after simultaneous exposure to odorant and PBS. The odorant solution consisted of 1 ml of 100 μM cadaverine (Sigma-Aldrich, St. Louis, MO) in PBS, whereas the control consisted of PBS. Each fish underwent three to four trials on the same day. Following each trial, fish were placed in an alternate experimental tank to acclimate for 60 minutes before commencing the next trial. The side of the odorant delivery was kept random for each trial.

### Scoring of olfactory-mediated behaviors

To analyze behavioral responses to cadaverine, we used recordings of the trials described above and manually coded behaviors. The cumulative durations of darting, freezing, and avoidance behaviors were quantified (in seconds) during the pre-odor and odor-exposure periods. Two or three investigators independently coded recordings, and their scores were averaged. We first analyzed darting behaviors, defined as rapid, high-velocity escape-like movements characterized by abrupt acceleration and directional changes. We also assessed freezing; a behavior defined as the absence of body displacement for ≥1 second. To assess cadaverine-evoked spatial avoidance, the arena was divided into two equal halves to define odor and non-odor halves during analysis. Spatial avoidance was quantified by measuring the cumulative time each fish spent in the odor and non-odor halves of the arena. Fish were considered to occupy a given half when the head crossed the midline. An odor avoidance index was calculated as: Odor avoidance Index = (T non-odor -T odor)/(Total time). Positive values indicate avoidance of cadaverine.

### Locomotor and exploratory assays in rectangular behavioral arena

We assessed the effects of hypoxia on locomotor ability and exploratory behaviors using the rectangular arena described above, since we aimed to assess swimming capabilities in the tanks used for odorant-evoked behavioral responses. We fasted and acclimated individual fish as described above, recorded their behavior for 30 seconds in the absence of cadaverine, and then analyzed the swimming and exploratory parameters in the video recordings.

### Analysis of locomotion and exploratory parameters in rectangular behavioral arena

We first measured vertical displacement by dividing the arena into three equal vertical zones during analysis. We calculated the cumulative vertical distance traveled by counting the number of crosses across the three zones and multiplying them by zone height (in cm). To assess overall exploratory activity in the rectangular arena, the tank was divided into four equal quadrants. Exploratory activity was quantified as the total number of vertical and horizontal quadrant boundary crossings per trial.

### Locomotor and exploratory assays in behavioral circular arena

To assess the effects of hypoxia on free-swimming parameters such as swimming speed, distance, and exploration rate in a larger tank, we used a circular arena, consisting of a 30-cm-diameter cylindrical behavioral tank filled with 2.5 liters of fish water. Fish were fasted and acclimated in similar circular tanks before trials. We placed an overhead digital camera 1 m above the arena to record behavior for 30 seconds in the absence of cadaverine exposure, and then analyzed swimming and exploratory parameters in the video recordings.

### Analysis of locomotion and exploratory parameters in circular behavioral arena

We used ToxTrac v024.1.1 (Rodriguez et al., 2018), an automated tracking software, to analyze behavior video recordings. We obtained the exploration rate from the software to quantify overall exploratory activity. Swimming distance (in cm) and speed (cm/sec) were also obtained from ToxTrac.

### Tissue processing

Control fish and hypoxic fish after recovery were sacrificed by over anesthetization with a solution of 0.03% tricaine (Sigma-Aldrich, St. Louis, MO) and decapitated. Whole heads were fixed with 4% paraformaldehyde (PFA; Sigma-Aldrich, St. Louis, MO) in PBS overnight (o/n) at 4° C. The next day, brains, olfactory organs, or whole heads were dissected and treated for immunohistochemistry (IHC) or histological staining.

Fixed brains and olfactory organs were washed and placed in 70% ethanol. Tissue was slowly dehydrated by incubations with increasing ethanol concentrations, followed by an incubation in xylene (Sigma-Aldrich, St. Louis, MO). We embedded dehydrated tissue by placing it in paraffin (Paraplast, St. Louis, MO) and rapidly cooling to solidification o/n. Then, we obtained semi-serial 10 μM sections of olfactory organs or brains in the coronal plane. These were adhered to charged slides (ThermoFisher, Waltham, MA) and left to dry o/n at 37° C. For whole-head processing, after fixing in PFA, tissue was decalcified using an RDO rapid decalcifying solution (Electron Microscopy Sciences, Hatfield, PA, USA) for 2 hrs at RT, washed with PBS, and placed in 70% ethanol. Tissue was dehydrated and embedded as described above, and then semi-serial, 10-μm sections in the horizontal plane were obtained and adhered to charged slides.

### Immunohistochemistry

Mounted tissue was rehydrated by descending ethanol incubations and then subjected to antigen retrieval with a 10 mM sodium citrate (Sigma Aldrich, Canada) solution (pH = 6.0) at 70° C for 10 minutes. Next, slides were washed with PBS and blocked for at least an hour with a blocking buffer containing 3% normal goat serum (NGS, Vector Laboratories, Burlingame, CA) and 0.4% Triton X-100 (Sigma Aldrich, St. Louis, MO). Following blocking, we incubated slides with primary antibodies (Table 1) o/n at RT and then incubated them with a fluorescently-labeled secondary antibody (Table 1) with 1 μg/ml of 4′,6-Diamidino-2-phenylindole (DAPI; BD Pharmingen) as a nuclear counterstain. Tissue sections were washed, coverslipped using PVA-DABCO (Sigma Aldrich, St. Louis, MO), and examined with a confocal laser-scanning microscope Nikon A1 and the NIS-Elements software (Nikon, Japan).

**Table 1.** Antibodies used in the study.

| Antibody | Species | Dilution | Source |
| --- | --- | --- | --- |
| HuC/D | Mouse | 1:100 | ThermoFisher (Waltham, MA) |
| GFAP | Rabbit | 1:500 | DAKO (Glostrup, Denmark) |
| Lcp1 | Rabbit | 1:100 | Genetex (Irvine, CA) |
| PCNA | Mouse | 1:1000 | Sigma Aldrich (St. Louis, MO) |
| Fluorescently-labeled secondaries: anti -mouse and -rabbit, IgG | Goat | 1:200 | Invitrogen (Waltham, MA) |

### Histological stainings and goblet cell quantification

Sectioned OE samples were deparaffinized with xylene, rehydrated by descending ethanol incubations, and stained with Alcian Blue and nuclear fast red (Vector Laboratories, Newark, CA). Then, we dehydrated the samples and mounted them using Permount (Fisher Scientific). Stained tissue was examined under light microscopy with Leica DM5000 B Automated Upright Microscope and analyzed with the Application Suite X System software (Leica, Wetzlar, Germany).

We identified goblet cells by their staining with Alcian blue, rounded morphology, and location within the olfactory lamellae. We manually quantified the number of glands per single olfactory lamellae in images taken at 40x magnification. We counted and averaged 4 to 6 sections per animal. To measure olfactory lamellae thickness and goblet cell diameter (in μm) in stained tissue, we used the Application Suite X System software (Leica, Wetzlar, Germany).

### Cell quantification and densitometry

We used Adobe Photoshop (Adobe, Mountain View, CA, USA) to quantify cells and to determine the optical density (OD) of immunostained tissue in images taken at 20× magnification. We quantified and averaged 3 to 5 tissue sections per animal. For cell quantification, we manually counted Lcp1+ and PCNA+ somata in OE and coronal telencephalon sections. We obtained OD measurements of immunostainings against Lcp1 and GFAP in OE and OB tissue, respectively, by converting images to 8-bit grayscale to obtain whole-section mean luminosity values, which were then converted to OD using the following formula: OD = −log (intensity of background/intensity of area of interest).

### TTC reduction assay

We evaluated brain mitochondrial dehydrogenase activity by using a 2,3,5-triphenyltetrazolium chloride (TTC) reduction assay. Immediately following fish decapitation, we dissected whole brains and incubated them in a 2% TTC (Sigma Aldrich, St. Louis, MO, USA) solution in PBS at 37 °C for 40 min, shielded from light. Then, TTC was removed, and 10% PFA was added to terminate the reaction. After 10 minutes, brains were washed in PBS, fixed in 4% PFA, and kept at 4 ° C o/n. The next day, we imaged the dorsal and ventral sides of TTC-stained brains using an Olympus SZ61 stereo microscope with a digital EP50 camera (Olympus, Waltham, MA), and then converted the images to 8-bit gray scale using Photoshop (Adobe, Mountain View, CA). We obtained the mean luminosity values for the dorsal and ventral sides of the OB and the rest of the brain. Luminosity values were converted to optical density (OD) by the following formula: OD = -log (intensity of background/intensity of area of interest). We calculated the percentage change using the formula: percentage change = (OD sample − mean OD control) / mean OD control value × 100.

Percent change values were calculated relative to control and multiplied by −1 so that negative values (more luminosity due to less TTC staining) indicate reduced mitochondrial activity.

### Statistical Analysis

Comparisons between groups were carried out using Analysis of Variance (ANOVA) with Tukey and Bonferroni’s post hoc tests, or unpaired t-tests. P-values less than 0.05 were considered significant. We used GraphPad Prism 10 software for all statistical analyses and graphs (GraphPad, San Diego, CA).

